# Olfactory ensheathing cells from the nasal mucosa and olfactory bulb have distinct membrane properties

**DOI:** 10.1101/640839

**Authors:** Katie E. Smith, Katherine Whitcroft, Stuart Law, Peter Andrews, David Choi, Daniel J. Jagger

## Abstract

Transplantation of Olfactory Ensheathing Cells (OECs) is a potential therapy for the regeneration of damaged neurons. While they maintain tissue homeostasis in the olfactory mucosa (OM) and olfactory bulb (OB), their regenerative properties also support the normal sense of smell by enabling continual turnover and axonal regrowth of olfactory sensory neurons (OSNs). However, the molecular physiology of OECs is not fully understood, especially that of OECs from the mucosa. Here, we carried out whole-cell patch clamp recordings from individual OECs cultured from the OM and OB of the adult rat, and from the human OM. A subset of OECs from the rat OM cultured 1-3 days in vitro (DIV) had large weakly rectifying K^+^ currents, which were sensitive to Ba^2+^ and desipramine, blockers of Kir4-family channels. Kir4.1 immunofluorescence was detectable in OM cells co-labelled for the OEC marker S100, and found adjacent to axons of OSNs. OECs cultured from rat OB had distinct properties though, displaying strongly rectifying inward currents at hyperpolarized membrane potentials and strongly rectifying outward currents at depolarized potentials. Kir4.1 immunofluorescence was not evident in OECs adjacent to axons of OSNs in the OB. A subset of human OECs cultured from the OM of adults had membrane properties comparable to those of the rat OM, i.e. dominated by Ba^2+^-sensitive weak inwardly rectifying currents. The membrane properties of peripheral OECs are different to those in central OECs, suggesting they may play distinct roles during olfaction.

**Table of Contents Image:** 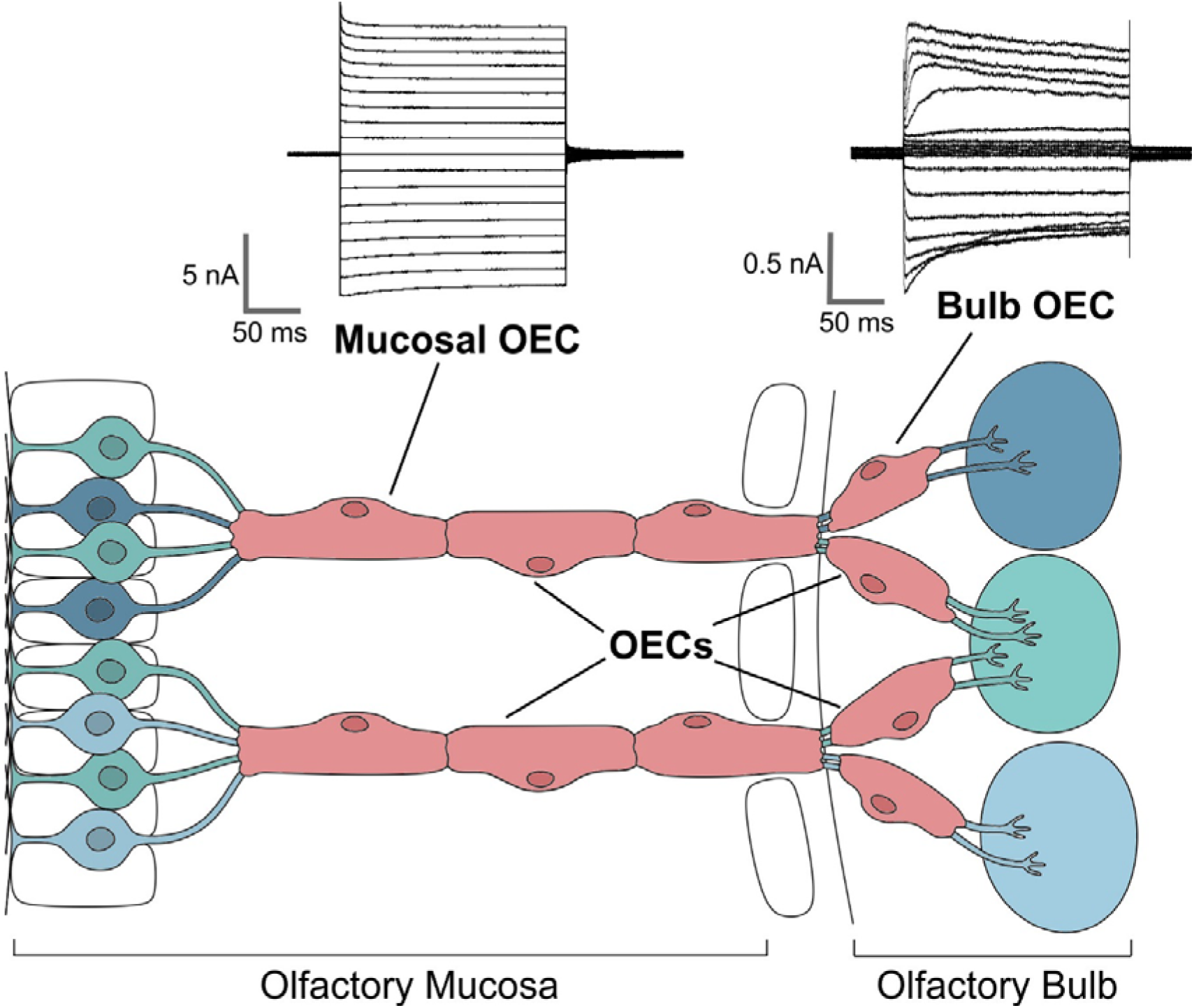

**Main points:**

- Peripheral and central OECs are functionally distinct
- Peripheral OECs have large weak inward rectifier currents
- Central OECs have strong inward and outward rectifier currents

## Introduction

Olfactory ensheathing cells (OECs) are specialized glia that bridge the boundary between the peripheral and central tissues mediating the sense of smell. The physiology of OECs remains poorly understood, but their inherent abilities appear to support the continual renewal of olfactory sensory neurons (OSNs) under normal conditions (Gomez et al., 2018). Around five to six million OSNs reside within the nasal sensory epithelium (de Castro, 2009; Jafek, 1983). Accompanied by OECs, their axons fasciculate to form the first cranial nerve that projects to the olfactory bulb (OB). These axons, which are thin and unmyelinated (Garcia-Gonzalez, Murcia-Belmonte, Clemente, & De Castro, 2013), enter the OB within the outer nerve layer (ONL), where they continue to be wrapped by OECs. The axons then coalesce into large glomeruli where they relay with mitral and tufted cells. The axons of these projection neurons form the lateral olfactory tract (LOT) that leads to the olfactory cortex where information is processed. The majority of axons within the LOT are myelinated, a process mediated by oligodendrocytes (Garcia-Gonzalez et al., 2013).

The lifespan of OSNs is relatively short, around 30 days in young mammals (Brann & Firestein, 2014), and so there is a need for continual neurogenesis in order to maintain a normal sense of smell. Harnessing the unusual neuro-regenerative abilities of the olfactory system continues to be promoted as a therapeutic approach to the reversal of spinal cord injury and certain demyelinating conditions (Assinck, Duncan, Hilton, Plemel, & Tetzlaff, 2017; Ekberg & St John, 2014; Gladwin & Choi, 2015; Gomez et al., 2018; Yao et al., 2018). The olfactory mucosa (OM) represents a particularly attractive target for sourcing OECs for clinical transplantation, as it permits relatively non-invasive endoscopic harvesting from affected patients, healthy volunteers, or cadaveric specimens (Andrews, Poirrier, Lund, & Choi, 2016; Choi & Gladwin, 2015). In addition, several lines of evidence suggest OECs derived from the OM may be especially beneficial for transplantation therapies (Yao et al., 2018). The physiological functions of peripheral OECs remain poorly understood at a cellular and molecular level (Chen, Kachramanoglou, Li, Andrews, & Choi, 2014; Gomez et al., 2018), and so it is currently unclear why this subtype of OECs could more successfully encourage neuronal regeneration. Much of our current understanding of neuro-glial interactions comes from structural, molecular and functional studies of rodent OECs, from either the OM or OB (Guerout et al., 2010; Hayat, Wigley, & Robbins, 2003; Piantanida et al., 2019; Rash et al., 2005; Rela, Bordey, & Greer, 2010; Rela, Piantanida, Bordey, & Greer, 2015; Rieger, Deitmer, & Lohr, 2007). Studies of excised human OM tissues have also helped identify common markers *in vivo* (Choi, Law, Raisman, & Li, 2008; Liu et al., 2010; Oprych, Cotfas, & Choi, 2017).

There is evidence of morphological, genetic, and functional heterogeneity between sub-populations of OECs (Gomez et al., 2018). OECs from the OB segregate into distinct phenotypic populations when maintained *in vitro* (Hayat et al., 2003). Some are spindle-shaped and Schwann cell-like, whereas others maintain a flattened astrocyte-like morphology, and these cell types display differences in their expression of glial markers (Honore et al., 2012). OECs within the OB and OM have distinct gene expression profiles (Guerout et al., 2010), but it is currently unclear whether these genetically and spatially separate populations mediate common physiological mechanisms that ensure normal homeostasis and continual neuronal turnover. Previous studies in acute slices of the OB from juvenile mice have described a variable contribution of certain types of membrane channels to shaping the electrical responses of OECs (Piantanida et al., 2019; Rela et al., 2010; Rela et al., 2015). Following the loss of gap-junctional coupling, these OECs display heterogeneous membrane currents, with variable contributions from hyperpolarization-activated inward rectifier K^+^ currents and depolarization-activated outward rectifier K^+^ currents. However, to date the molecular identities of the ion channel subtypes underlying these responses in central OECs remain undetermined. In addition, there is a lack of information available on the membrane properties of peripheral OECs.

In this study, we employed whole-cell patch clamp to investigate the membrane properties of OECs cultured from the OM of adult rats, and found them to be distinct to those cultured from the rat OB. There was a strong resemblance, though, between the properties of rat OM OECs and of those harvested by endoscopic biopsy from the OM of humans. Weak rectifier currents dominated the electrical responses in OECs from the OM of both rats and humans, and consistent with these observations, there was expression of weak rectifier K^+^ channel subunits by OECs within lamina propria of the rat OM, and in cultured human OECs. The data provide the first description of the membrane properties of peripheral OECs, and highlight region-specific differences that may help explain the physiological roles of these enigmatic glial cells.

## Materials and Methods

### Animals

Twelve-week old Sprague Dawley rats of either sex (purchased from Charles River or Harlan) were killed by CO_2_ inhalation followed by cervical dislocation. All animal work conformed to United Kingdom legislation outlined in the Animals (Scientific Procedures) Act 1986.

### Human OEC donors

Adult patients undergoing functional septorhinoplasty under the care of PA were recruited. The surgical procedures for harvesting patient specimens largely followed those outlined elsewhere (Andrews et al., 2016; Kachramanoglou, Law, Andrews, Li, & Choi, 2013). All procedures were performed under NHS Research Ethics Committee approval (14/SC/1180) and patient informed consent. The patients’ medical history was documented including age, sex, presenting complaint, past medical and surgical history, drug history including recent use of topical and/or oral steroids, allergies, and smoking status. Birhinal olfactory function was assessed using “Sniffin’ Sticks”, a validated psychophysical tool which separately tests odor threshold, discrimination and identification (Hummel, Kobal, Gudziol, & Mackay-Sim, 2007; Neumann et al., 2012). Detailed testing procedures are outlined elsewhere (Whitcroft, Cuevas, Haehner, & Hummel, 2017). Normosmia was assumed where composite threshold, discrimination and identification (TDI) score was ≥30.3 (Hummel et al., 2007). Patients with neurological, psychiatric or other conditions known to affect olfaction were excluded, as were those under 18 or over 70 years of age. Samples from three participants were obtained (male:female = 2:1; mean age 21 years, range 18-29). Two of the participants were normosmic according composite TDI score. One participant scored within the high hyposmic range (TDI 27.75), though this was associated with significant nasal obstruction at the time of testing, and normalized at three months post-operatively.

### Surgical procedure for human olfactory biopsies

Biopsies were harvested endoscopically during functional septorhinoplasty. A detailed description of the technique is outlined elsewhere (Kachramanoglou et al., 2013). In brief, full thickness biopsies were obtained (including bone, so ensuring integrity of the lamina propria) from the middle third of the superior turbinate. The nasal mucosa was prepared either with topical lignocaine with phenylephrine (lignocaine hydrochloride 5% and phenylephrine hydrochloride 0.5%) or Moffett’s solution (2 mL 6% cocaine, 1 mL 1:1000 epinephrine, 2 mL 8% sodium bicarbonate solution), and the procedure performed under general anesthetic. Further vasoconstriction was obtained as required through use of topical epinephrine (1:10,000) soaked neuro patties. Once obtained, the specimen was immediately placed into ice-cold culture medium (DMEM/F-12 + GlutaMax, 10% fetal bovine serum (FBS), 1% penicillin-streptomycin, 1% insulin-transferrin-selenium) with 10% deactivated fetal calf serum (Invitrogen), and transferred to the laboratory.

### Rat and human OEC cultures

The olfactory tissue from two rats was used for each culture. For rat or human mucosal OEC cultures, mucosa were first washed in Hanks balanced salt solution (no Ca^2+^/Mg^2+^) to remove mucus before incubating in 2 ml dissociation media containing 2.4 U/ml Dispase II (Sigma) and 0.5% collagenase Type I (Sigma) in DMEM/F12 + GlutaMax (ThermoFisher) for 30-60 minutes at 37 °C. Following digestion, the tissue was triturated, first with a wide-bore fire-polished pipette (5x) then with a narrow-bore pipette (15-20x). Growth medium (DMEM/F-12 + GlutaMax, 10% FBS, 1% penicillin-streptomycin, 1% insulin-transferrin-selenium) was added, and the dissociated cells were centrifuged at 300 xg for 5 minutes. The supernatant was discarded and the cell pellet re-suspended in growth medium. The cell suspension was plated at varying densities (between 5,000 and 10,000 cells per coverslip) onto poly-L-lysine coated coverslips (100 *µ*g/ml), and then maintained in a humidified incubator at 37 °C and 5% CO_2_ for 1-3 days in vitro (DIV). For rat bulb OEC cultures, the two outermost layers of the olfactory bulb were digested in 1% trypsin (Worthington Biochemicals) in Hank’s balanced salt solution (no Ca^2+^/Mg^2+^) at 37 °C for 10 minutes. Growth medium was added to quench the reaction, and the tissue was then allowed to settle at the bottom of the tube. All but ∼1 ml of excess solution was removed and DNase I (400 U; ThermoFisher) added. The tissue was then triturated as described for the mucosal tissue. Growth media was added and the cells were centrifuged at 300 xg for 5 minutes. The supernatant was removed and the cell pellet re-suspended in growth medium. The cell suspension was then plated and incubated as described for the mucosal cultures.

### Electrophysiology

Whole-cell patch clamp recordings were performed at room temperature, using an Axopatch 200B amplifier and a Digidata board (Molecular Devices) under the control of pClamp acquisition software (version 8, Molecular Devices). Cells were super-fused continuously with an external solution containing (in mM): 145 NaCl, 4 KCl, 1 MgCl_2_, 1.3 CaCl_2_, 10 HEPES and 5 glucose, pH 7.3. BaCl_2_ and desipramine hydrochloride (Sigma-Aldrich) were prepared as 1 M and 100 mM stock solutions in water, respectively, before dilution in artificial perilymph solution to a final concentration of 100 *µ*M. Patch pipettes were pulled from borosilicate glass (GC120TF-10, Harvard Apparatus) and had a resistance of 3-6.5 MΩ when filled with intracellular solution containing (in mM): 130 K-gluconate, 5 KCl, 2 MgATP, 2 Na_2_ATP, 0.3 Na_3_GTP, 10 Na_2_-phosphocreatine, 1 EGTA and 10 HEPES, pH 7.2. Series resistance was compensated online by 70%. The liquid junction potential, measured at −13 mV, was subtracted offline. Resting membrane potential was measured using the *I=0* circuitry of the amplifier. The voltage clamp protocol used to assess the currents is described in the Results section and Figure Legends. To enable comparison of the current-voltage relationship of individual cells, whole-cell currents were normalized to the maximum inward current for each cell. All group data is expressed as mean ± standard error of the mean (SE). Data was obtained from four independent cultures for rat OM and rat OB experiments, and from three independent cultures for human OM experiments.

### Tissue sectioning and immunofluorescence

Rat OB and OM were fixed in 4% paraformaldehyde (PFA) for 1 hour at room temperature, and then mounted in 4% low-melting temperature agarose, before sectioning at 150 *µ*m intervals on a vibratome (1000 plus system, Intracel). Tissues were not decalcified prior to sectioning. Cells cultured on coverslips were fixed in 4% PFA for 20 minutes at room temperature, and then stored in PBS. All steps in the immunofluorescence protocol were performed at room temperature. Sections and cells were blocked and permeabilized in 10% normal goat serum and 0.2% Triton-X-100 in PBS for 1 hour, and then incubated in primary antibodies diluted in lysine blocking solution (0.1 M lysine and 0.2% Triton-X-100 in PBS) for 3 hours, followed by three 10 minute washes in PBS. When using primary antibodies raised in goat, normal goat serum was substituted by FBS. Sections or cells were incubated in the secondary antibodies diluted 1:400 in lysine blocking solution for 1 hour, followed by three further washes in PBS. For imaging, sections were mounted in glass bottom dishes (MatTek Corporation) containing Vectashield mounting medium with DAPI (Vector Laboratories), whereas cells on coverslips were mounted on glass slides. Confocal images were acquired with a laser scanning confocal microscope (Carl Zeiss) equipped with 10x (N.A. 0.3), 20x (N.A. 0.75), and 63x (N.A. 1.2) objectives. The images shown are maximum intensity z-projections of 3-9 adjacent z-planes, except Figures 3B and 3C that are single z-planes.

**Figure 1.**
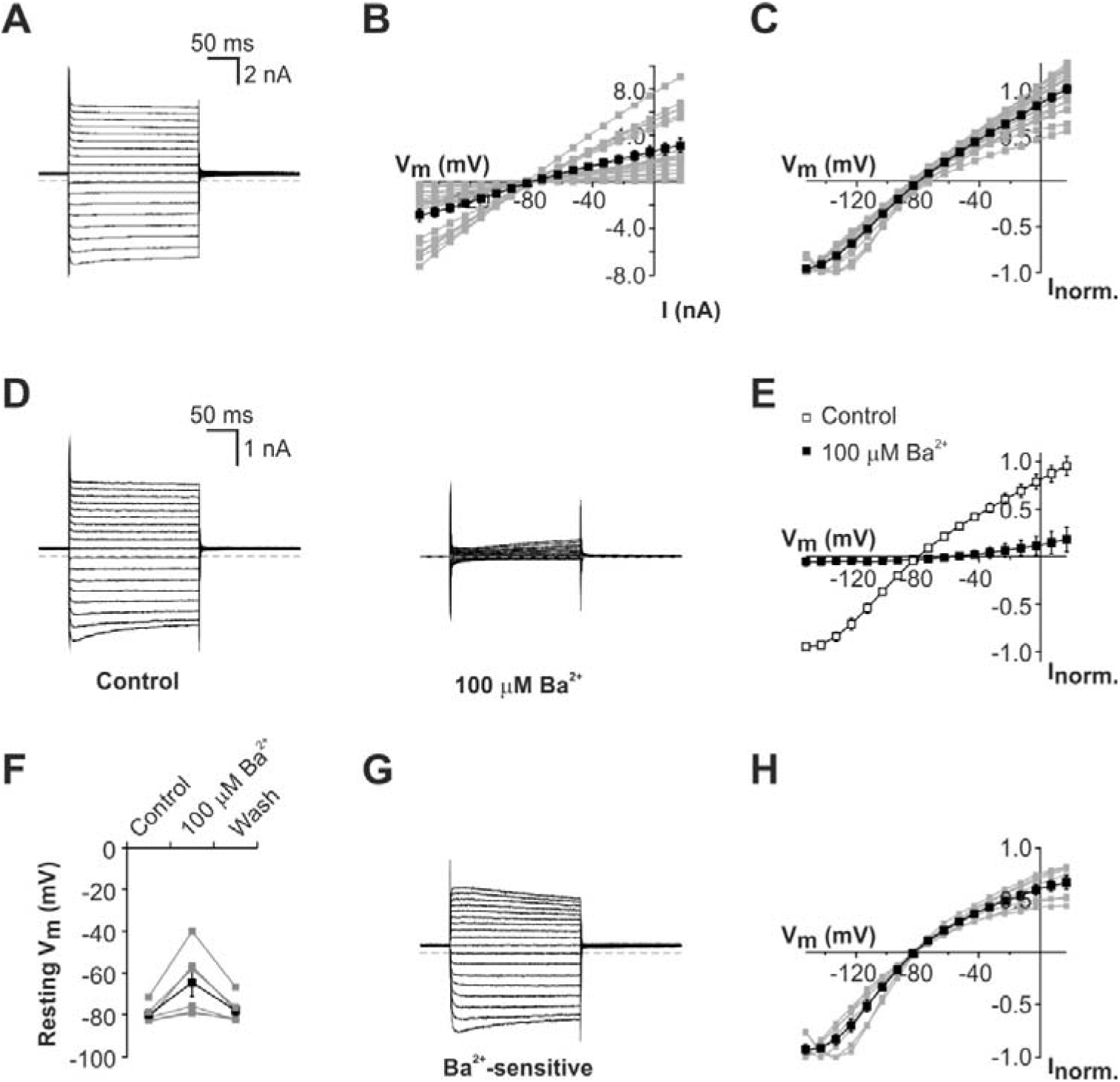
Weak rectifier currents in OECs cultured from the rat olfactory mucosa. ***A*** Whole-cell voltage clamp recording from an OEC following three days *in vitro* (3 DIV), showing membrane currents activated by sequential 200 ms voltage steps between −153 mV and 17 mV, in 10 mV increments from a holding potential of −73 mV. The discontinuous line denotes the position of zero-current. Hyperpolarizing steps activated large inward currents, whereas depolarizing steps typically activated outward currents. ***B*** Group (mean ± SE, black squares) and individual (gray squares) current-voltage (*I-V*) relationships of 19 OECs after 1-3 DIV. ***C*** Normalized group and individual *I-V* plots demonstrated the weak voltage-dependence of the whole-cell currents. ***D*** Currents in a representative OEC before (control) and after bath application of 100 *µ*M Ba^2+^. ***E*** Group *I-V* relationships before (open squares) and after (filled squares) Ba^2+^ application in seven OECs. ***F*** Bath applied Ba^2+^ depolarized the resting membrane potential (V_m_) of all cells reversibly (n = 6). ***G*** Digital subtraction of the currents in the presence of Ba^2+^ from control currents revealed a weak inwardly rectifying Ba^2+^-sensitive current. ***H*** Group (black squares) and individual (gray squares) *I-V* relationships of Ba^2+^-sensitive current in seven OECs.

**Figure 2.**
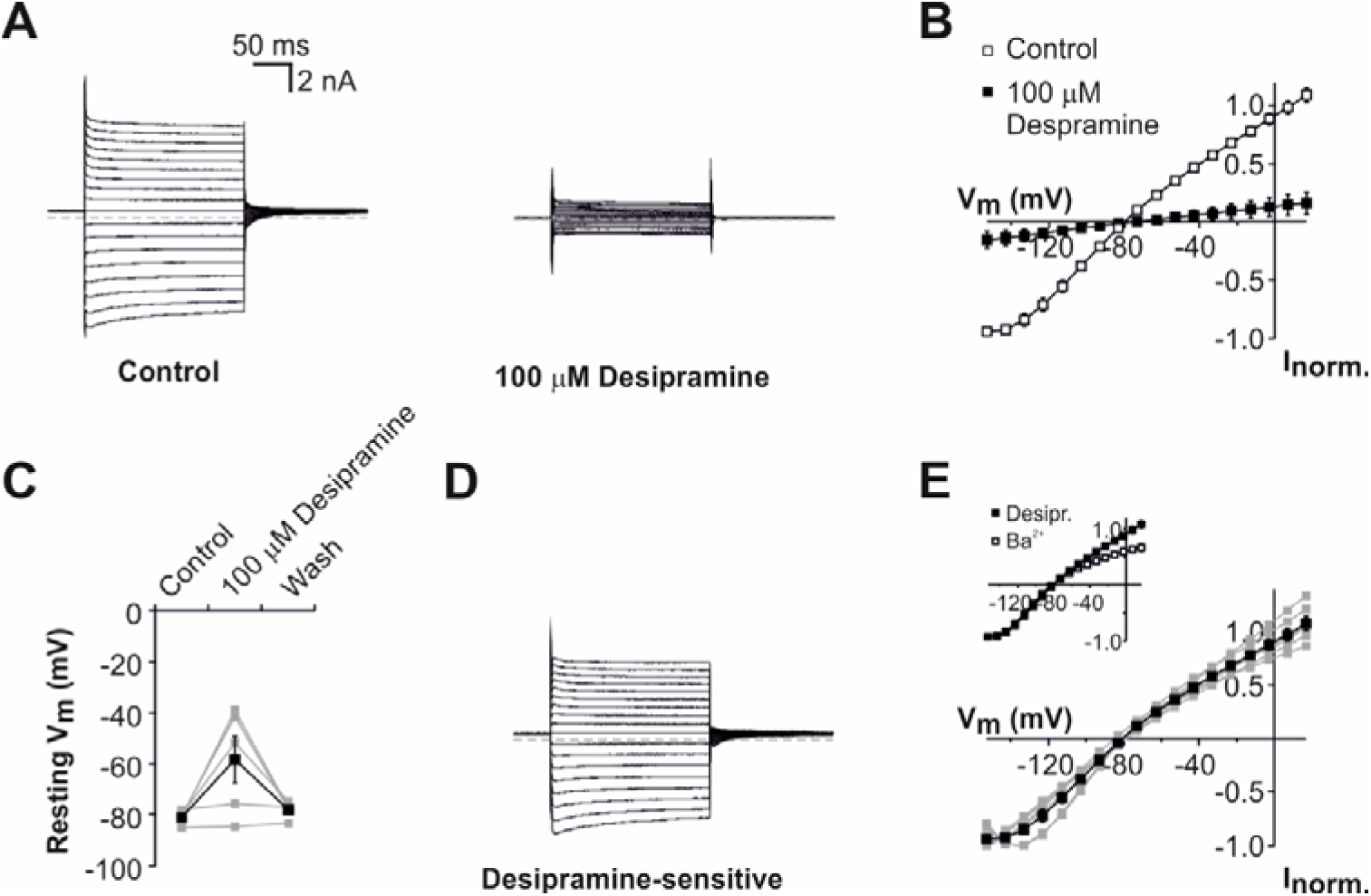
Weakly rectifying currents in rat olfactory mucosa OECs are sensitive to the Kir4 blocker desipramine. ***A*** Currents in a representative OEC before (control) and after bath application of 100 *µ*M desipramine. ***B*** Group *I-V* relationships before (open squares) and after (filled squares) desipramine application in six OECs. ***C*** Bath applied desipramine depolarized the resting membrane potential (V_m_) of 4/5 cells reversibly. ***D*** Digital subtraction of the currents in the presence of desipramine from control currents revealed a weak inwardly rectifying desipramine-sensitive current. ***E*** Group (black squares) and individual (gray squares) *I-V* relationships of desipramine-sensitive currents in six OECs. *Inset*, comparison of the Ba^2+^-sensitive and desipramine-sensitive components of membrane currents recorded in OECs from rat OM (group data shown in Fig 1H and Fig 2E, respectively).

**Figure 3.**
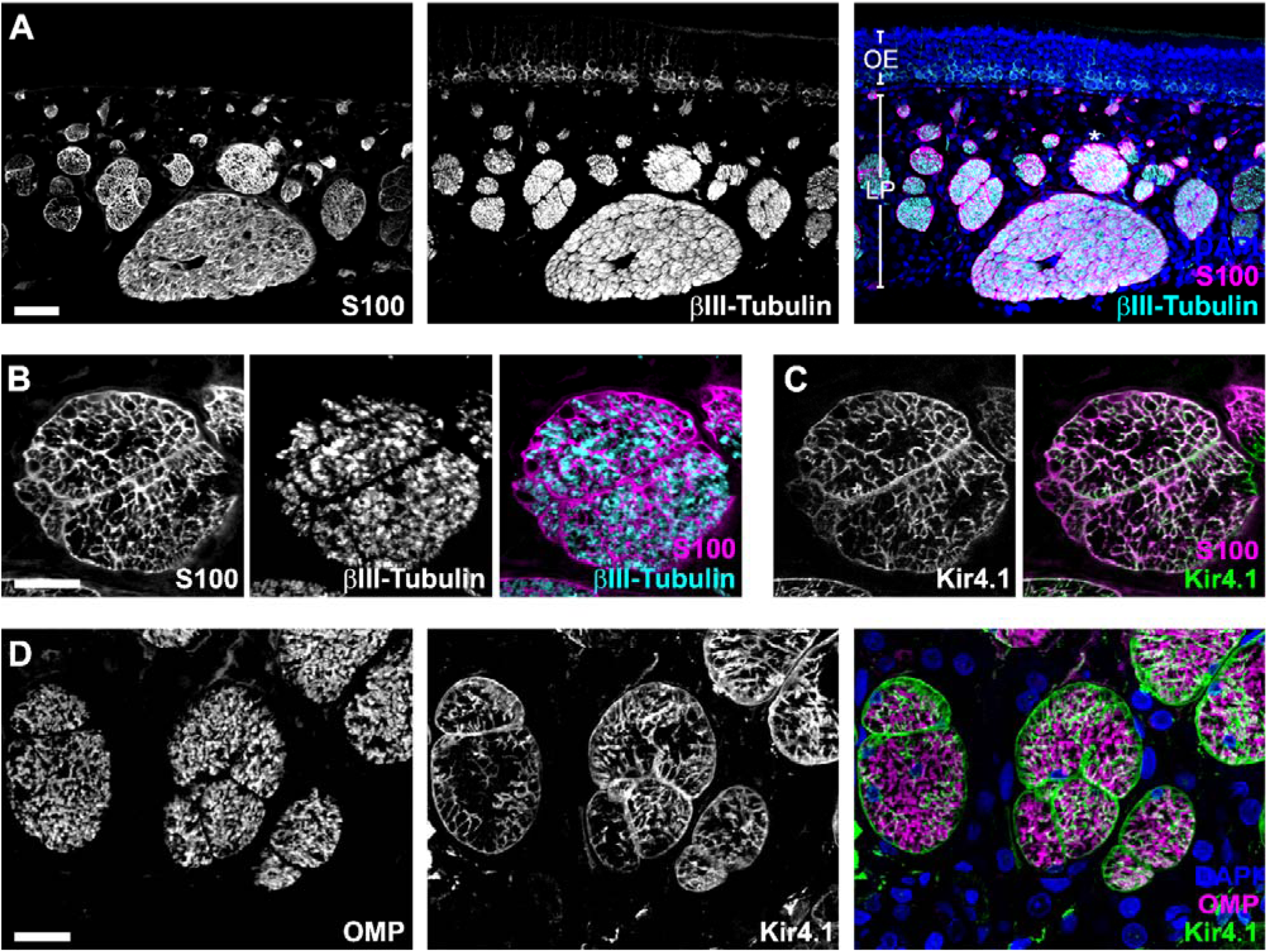
OECs in the rat olfactory mucosa express Kir4.1 channel subunits. ***A*** Horizontal section of rat OM, immuno-labelled for the OEC marker S100 (*left*) and the neuronal marker tubulin (*center*). The overlay has cell nuclei counter-stained using DAPI (*right*). The cell bodies of OSNs reside within the olfactory epithelium (OE), and their axons fasciculate within lamina propria (LP). S100+ OECs lie beneath the basement membrane and in close association with SN axons. **B** and **C** show detail of a fascicle highlighted in ***A*** (*), labelled for S100 and βIII-tubulin (***B***), and for the weakly rectifying K^+^ channel subunit Kir4.1 (***C***). ***D*** shows detail of fascicles where OSN axons are labelled using anti-OMP and OECs are labelled using anti-Kir4.1. Scale bars: ***A*** 50 *µ*m, ***B-D*** 20 *µ*m.

### Antibodies

The following primary antibodies were used for immunofluorescence: rabbit polyclonal anti-Kir4.1 (APC-035, Alomone Labs); guinea pig polyclonal anti-Kir4.1 (AGP-012, Alomone Labs); mouse monoclonal βIII-tubulin (subtype IgG_2a_; clone TUJ1; Covance); rabbit polyclonal anti-S100 (Z0311; Dako); Goat polyclonal anti-OMP (Wako). AlexaFluor-tagged secondary antibodies (Life Technologies) were used to detect primary antibodies: AlexaFluor-488 goat anti-guinea pig; AlexaFluor-633 goat anti-mouse IgG_2a_, AlexaFluor-488 donkey anti-mouse; AlexaFluor-555 donkey anti-rabbit; AlexaFluor-633 donkey anti-goat.

## Results

The main aim of this study was to determine the electrophysiological properties of cultured peripheral OECs. These experiments could enable us (1) to identify ion channel mechanisms that underlie their functions in vivo, and (2) identify protein markers that could help refine cell isolation procedures, and (3) to allow functional assessment of biopsied cells obtained for transplantation therapies. As a first part of this procedure, we studied the properties of cultured OECs from the rat OM and then OECs from the rat OB, which provided a first comparative analysis of these cells obtained from different regions. These experiments allowed us to determine the best conditions for effective cell isolations, and that our experimental conditions could maintain OECs appropriately for patch clamp recordings. With these conditions, we could then efficiently characterize the properties of OECs within valuable human OM biopsies.

### OECs cultured from the rat olfactory mucosa display weak inwardly rectifying membrane currents

Using approaches previously described for human cells (Kachramanoglou et al., 2013), we prepared dissociated cell cultures of OECs harvested from the rat OM. Whole-cell patch clamp experiments were then carried out following 1-3 DIV. Given the possibility of intercellular gap junctional coupling, as reported for OECs in mouse OB (Rela et al., 2010), our recordings targeted isolated cells. In response to a series of 200 ms voltage steps, a prominent subset of mucosal OECS (19/42 cells) displayed weak inwardly rectifying or non-rectifying currents (**Figure 1A-C**). Cells with these current profiles had hyperpolarized resting membrane potentials of −79.3 ± 0.9 mV (n = 19). 100 *µ*M Ba^2+^ was applied to the bath solution to determine whether Kir channels mediated the weak inwardly rectifying currents, (**Figure 1D,E**). Ba^2+^ application resulted in a substantial block of the membrane currents (**Figure 1D,E**; control current amplitude at −153 mV = −3.28 ± 1.04 nA, Ba^2+^ current amplitude = −0.20 ± 0.09 nA, n=7). Ba^2+^ application caused a modest depolarization of the resting membrane potential that was reversible on washout (**Figure 1F**; control V_m_ = −79.80 ± 1.81 mV, Ba^2+^ V_m_ = −64.65 ± 6.43 mV, wash V_m_ = −77.85 ± 2.46 mV, n=6). The digitally subtracted Ba^2+^-sensitive currents displayed weak inward rectification (**Figure 1G,H**). Accordingly, the weak inwardly rectifying currents were also sensitive to the tricyclic anti-depressant desipramine (100 *µ*M; **Figure 2A,B**; control current amplitude at −153 mV = −3.74 ± 1.02 nA, desipramine current amplitude = −0.48 ± 0.21 nA, n=6), a known blocker of Kir4-family channels (Su et al., 2007). Like Ba^2+^, desipramine had a depolarizing effect on the resting membrane potential of the cells, which was reversible on washout (**Figure 2C**; control V_m_ = −81.06 ± 1.16 mV, desipramine V_m_ = −58.42 ± 9.18 mV, wash V_m_ = −78.02 ± 1.39 mV, n=5). The kinetics and voltage-dependence of the digitally subtracted desipramine-sensitive whole-cell currents bore a strong resemblance to those blocked by Ba^2+^ (**Figure 2D,E**).

Prompted by the similarities between these currents and those in other glial cells mediated by Kir4.1 channel subunits (Butt & Kalsi, 2006; Seifert, Henneberger, & Steinhauser, 2018; Tang, Schmidt, Perez-Leighton, & Kofuji, 2010), we next used immunofluorescence to determine whether these subunits were expressed within OM tissue. In sections cut from the rat OM, S100+ OECs were in close association with OSN axons running through lamina propria (**Figure 3A,B**). The Kir4.1 antibody also labelled the S100+ OECs (**Figure 3C**). Within each fasciculation, Kir4.1+ OECs surrounded axons labelled by the olfactory sensory neuron marker OMP (**Figure 3D**).

### The membrane properties of peripheral and central OECs are different

Previous studies in the mouse OB have reported that once gap junction channels are blocked, the OECs have non-linear current profiles (Rela et al., 2010; Rela et al., 2015). These responses point to variable contributions from strong inward rectifier and strong outward rectifier currents, distinct from the weak rectification profile we observed in OECs from the rat OM. To investigate possible differences in membrane properties of peripheral and central OECs, we next prepared cultures from the rat OB. In 16/23 cells (1-3 DIV) there was a strong inward rectification in response to hyperpolarizing voltage steps below −80 mV, and a strong outward rectification in response to depolarizing voltage steps above −40 mV (**Figure 4A-C**). In the presence of Ba^2+^, only strong outwardly rectifying currents remained (**Figure 4D,E**). The digitally subtracted Ba^2+^-sensitive inward currents displayed a strongly rectifying profile (**Figure 4F,G**), with little or no outward component at depolarized potentials (**Figure 4G**). Direct comparison of the mean Ba^2+^-sensitive currents in OECs derived from OB and OM clearly highlighted their contrasting properties (**Figure 4H**).

**Figure 4.**
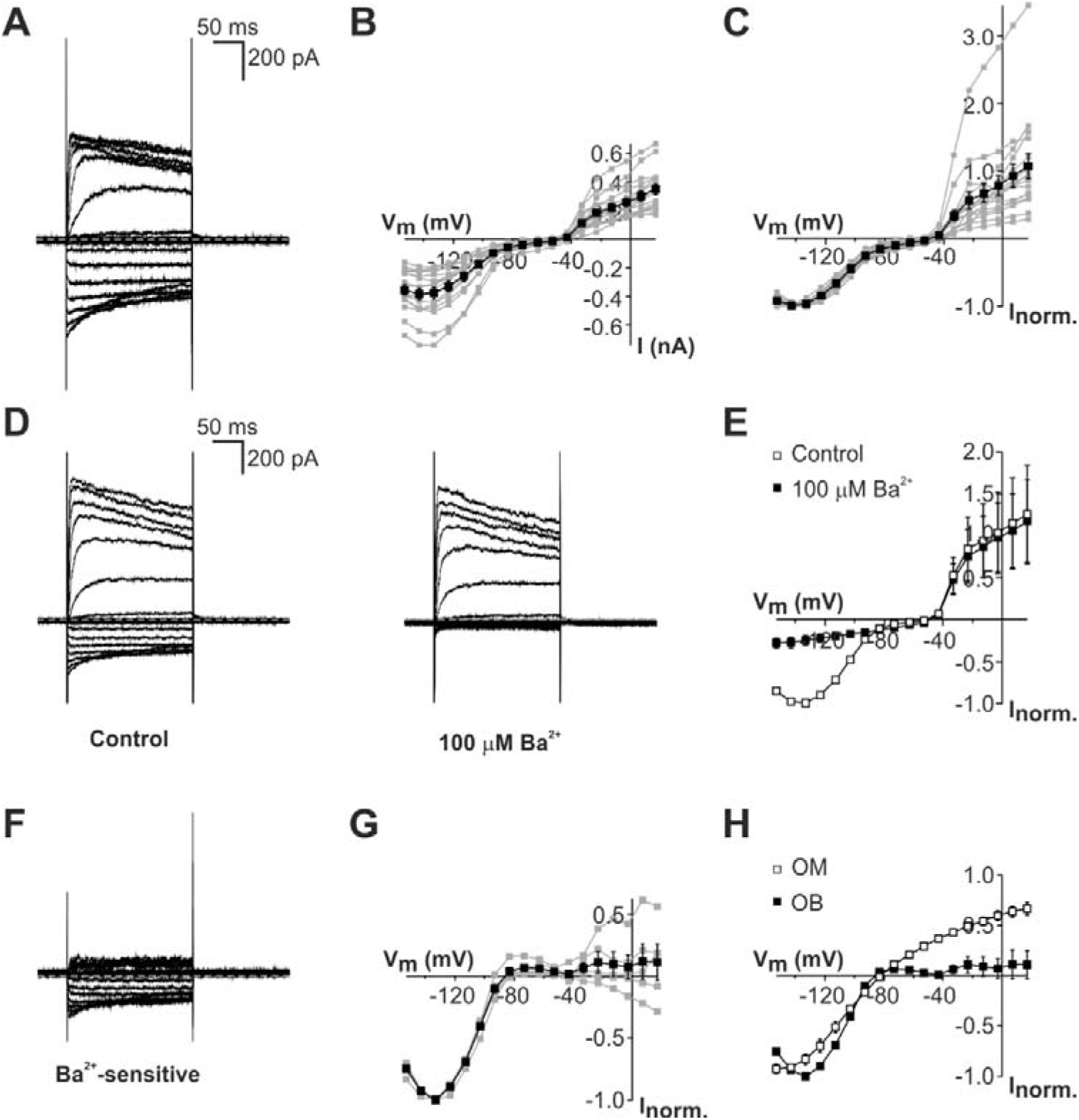
OECs cultured from the rat olfactory bulb have membrane properties distinct to OECs from the rat olfactory mucosa. ***A*** Whole-cell voltage clamp recording from a representative OEC following three days *in vitro* (3 DIV), showing membrane currents activated by sequential 200 ms voltage steps between −153 mV and 17 mV, in 10 mV increments from a holding potential of −73 mV. Hyperpolarizing steps below around −80 mV activated inward currents, whereas depolarizing steps above −40 mV typically activated smaller outward currents. ***B*** Group (mean ± SE, black squares) and individual (gray squares) current-voltage (*I-V*) relationships of 16 OECs at 1-3 DIV. ***C*** Normalized *I-V* plots demonstrated the pronounced voltage-dependence of both inward and outward currents. ***D*** Currents before (control) and after bath application of 100 *µ*M Ba^2+^ in a representative OEC. ***E*** Group *I-V* relationships before (open squares) and after (filled squares) Ba^2+^ application (n = 5). ***F*** Digital subtraction of the currents in the presence of Ba^2+^ from control currents revealed a strong inwardly rectifying Ba^2+^-sensitive current. ***G*** Group (black squares) and individual (gray squares) *I-V* relationships of Ba^2+^-sensitive current in five OECs. ***H*** Comparison of the Ba^2+^-sensitive component of membrane currents recorded in OECs from rat OB and OM (group data shown in Fig 4G and Fig 1H, respectively).

Given the apparent lack of weak rectifier currents in central OECs, we next carried out immunofluorescence using sections of rat OB. S100+ OECs were evident in close association with OSN axons within the ONL, and within the GL (**Figure 5A**). In agreement with previous observations in mice (Rela et al., 2015), Kir4.1 immuno-reactivity was largely absent from OECs within the ONL (**Figure 5A,B**). Diffuse Kir4.1 immunofluorescence was evident in S100+ cells within the glomeruli though (**Figure 5A**), and within large multipolar cells at the border of the ONL and GL (**Figure 5B**). Together, these experiments demonstrate that ion channel expression in OECs is regionally specific, an observation that suggests these cells play different roles in the peripheral and central nervous tissue environments.

**Figure 5.**
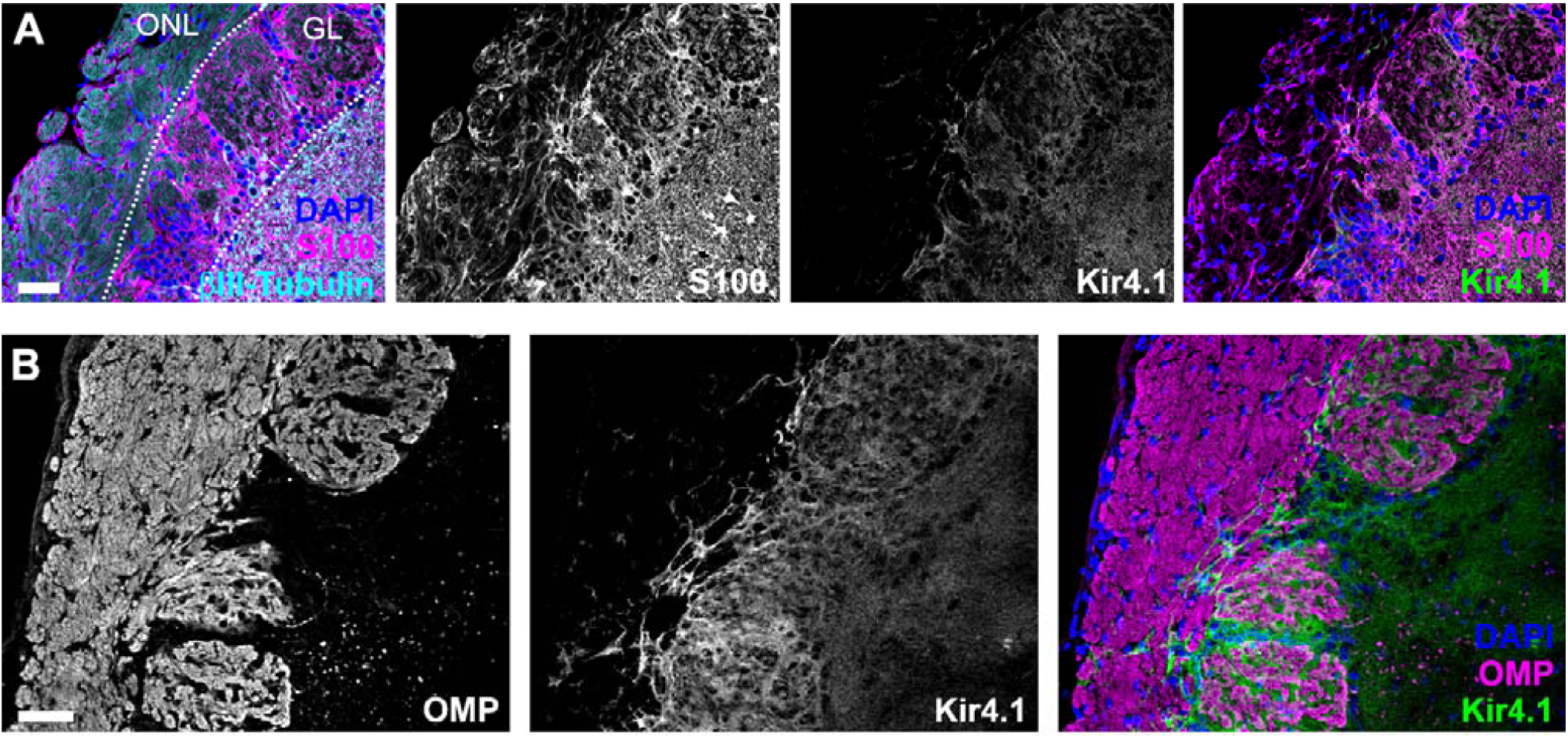
OECs in the outer olfactory nerve layer of the rat olfactory bulb lack expression of Kir4.1. ***A*** Sagittal section of rat OB showing OSN axons (labelled for the neuronal marker βIII-tubulin) running through the outer neuronal layer (ONL), before entering the glomerular layer (GL). Glial cells, including OECs in the ONL, are immuno-labelled for S100. Kir4.1 immunofluorescence was largely undetectable in the ONL, but was evident in cells of the deeper layers. The overlays have cell nuclei counter-stained using DAPI. ***B*** A different sagittal section of rat OB, immuno-labelled for the OSN marker OMP and Kir4.1. Kir4.1 immunofluorescence was largely undetectable in the ONL, and was restricted to large multi-polar cells within the GL. The overlay has cell nuclei counter-stained using DAPI. Scale bars: 50 *µ*m.

### OECs cultured from the rat and human olfactory mucosa are functionally comparable

Having validated our culturing and recording conditions using rat cells, in a final group of experiments we aimed to determine the electrophysiological profile of OECs harvested by endoscopic biopsies of human OM. In response to the same voltage step protocol applied to rat mucosal OECs (e.g. **Figure 1A**), a prominent subpopulation of human OECs (11/29, 1-3 DIV) displayed weak inwardly rectifying currents (**Figure 6A-C**). Cells with this type of current had hyperpolarized resting membrane potentials of −80.2 ± 1.0 mV (n = 11). 100 *µ*M Ba^2+^ application resulted in a substantial block of the weak inwardly rectifying current (**Figure 6D,E**; control current amplitude at −153 mV = −1.32 ± 0.2 nA, Ba^2+^ current amplitude = −0.16 ± 0.04 nA, n=6). Ba^2+^ also caused a depolarization of the resting membrane potential of human OECs, which was reversible upon washout (**Figure 6F**; control V_m_ = −81.42 ± 1.36 mV, n=6; Ba^2+^ V_m_ = −54.63 ± 3.74 mV, n=6; wash V_m_ = −79.08 ± 1.32 mV, n=5). The digitally subtracted Ba^2+^-sensitive currents showed weak inward rectification (**Figure 6G,H**) and direct comparison of the Ba^2+^-sensitive currents of OECs from human and rat OM revealed an almost complete overlap in their current-voltage relationship (**Figure 6H** *inset*). We did not identify Kir4.1+ OECs in fixed sections prepared from biopsies of human OM (not shown), but there was a sub-population of cells in the human OM cultures which were double labelled for S100 and Kir4.1 (**Figure 6I**) suggesting that Kir4.1 channels mediate the weak inwardly rectifying currents.

**Figure 6.**
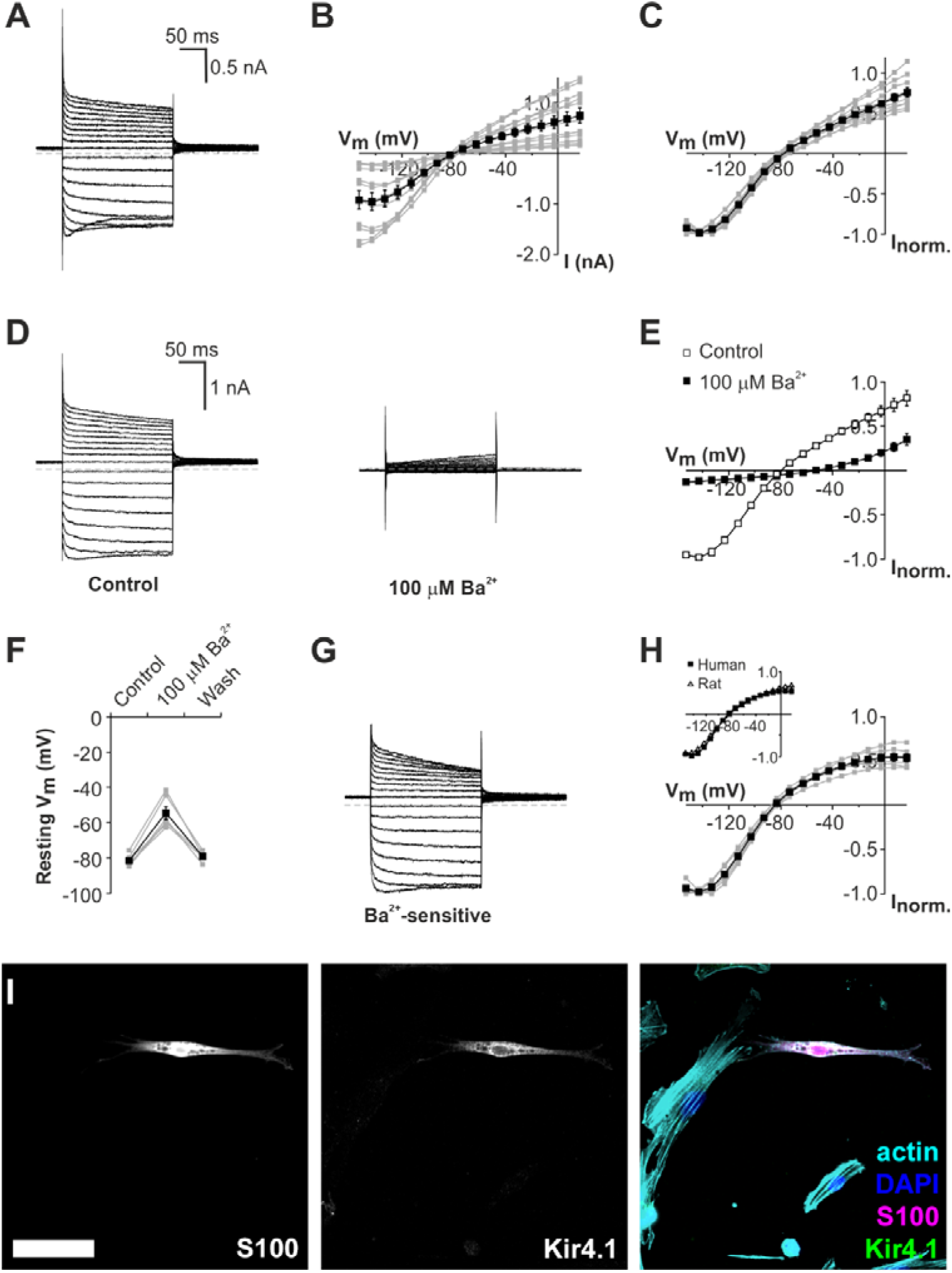
Membrane properties of OECs from the human olfactory mucosa are comparable to those from the rat olfactory mucosa. ***A*** Whole-cell voltage clamp recording from a representative human OEC following three days *in vitro* (3 DIV), showing membrane currents activated by sequential 200 ms voltage steps between −153 mV and 17 mV, in 10 mV increments from a holding potential of −73 mV. Hyperpolarizing steps activated large inward currents, whereas depolarizing steps typically activated outward currents. ***B*** Group (black squares) and individual (gray squares) *I-V* relationships of 11 human OECs after 1-3 DIV. ***C*** Normalized group and individual *I-V* plots demonstrated the weak voltage-dependence of the whole-cell currents. ***D*** Currents in a representative human OEC before (control) and after bath application of 100 *µ*M Ba^2+^. ***E*** Group *I-V* relationships before (open squares) and after (filled squares) Ba^2+^ application in 6 OECs. ***F*** Bath applied Ba^2+^ depolarized the resting membrane potential (V_m_) of all cells reversibly. ***G*** Digital subtraction of the currents in the presence of Ba^2+^ from control currents revealed a weak inwardly rectifying Ba^2+^-sensitive current. ***H*** Group (black squares) and individual (gray squares) *I-V* relationships of Ba^2+^-sensitive currents in six OECs. *Inset*, comparison of the Ba^2+^-sensitive component of whole-cell currents recorded in OECs from human OM and rat OM (group data shown in Fig 6H and Fig 1H, respectively). ***I*** In a 3 DIV culture, an OEC co-labeled with anti-S100 (*left* panel) and anti-Kir4.1 antibodies (*center*). The overlay (*right*) has cell nuclei counter-stained using DAPI (blue), and F-actin using phalloidin (cyan). Scale bar: 50 *µ*m.

## Discussion

The main findings of this study are (1) that there are differences in the membrane physiology of OECs derived from the olfactory periphery and those that reside within the central nervous system (CNS), and (2) that OECs biopsied from the OM display membrane properties that are consistent with roles in K^+^ homeostasis during the transduction of olfactory stimuli. These results should improve our understanding of the physiological roles played by these specialized glial cells during normal olfaction, and the description of distinct membrane properties within peripheral and central OECs may help explain how the two types of OECs behave when transplanted into the damaged or diseased CNS.

Our experiments reveal previously unrecognized qualities of OECs resident in the olfactory periphery. Whole-cell patch clamp recordings identified weak inward rectifier currents that were sensitive to both Ba^2+^ and desipramine, indicating a likely involvement of Kir4-family channel subunits. Ba^2+^-sensitive currents in rat and human OECs from the OM had near-identical kinetics and voltage-dependence. The reversible depolarization of the resting membrane potential during pharmacological block suggests that these currents contribute to the hyperpolarised resting membrane potential of OECs. A similar role for weak inwardly rectifying channels has been proposed for astrocytes (Butt & Kalsi, 2006) and satellite glial cells (Tang et al., 2010). The expression of these channels in glia has been associated with the maintenance of physiological levels of K^+^ within the extracellular space, particularly during periods of high neuronal activity in the CNS (Bay & Butt, 2012; Butt & Kalsi, 2006; Higashi et al., 2001). Satellite glial cells in peripheral ganglia also express these weakly rectifying channels at high densities, suggesting comparable homeostatic roles support afferent signaling within various sensory modalities (Hibino et al., 1999; Tang et al., 2010).

The patch clamp data prompted our subsequent detection of Kir4.1 channel subunits within rat OECs lying in close association with the unmyelinated axons of OSNs. S100+/Kir4.1+ OECs were evident within the centrally directed axonal bundles running within lamina propria. Unfortunately, in the present study we were unable to obtain consistent Kir4.1 immunofluorescence from sections derived from the human OM. The reasons for this are currently unclear, but this may have been due to limitations of the fixation achievable in the biopsied human tissue. However, consistent with the electrophysiology experiments on human OM cultures, we were able to identify cultured S100+ cells that were also immuno-positive for Kir4.1.

In agreement with an earlier study in mice (Rela et al., 2015), we did not detect Kir4.1 immunofluorescence within the ONL of the rat OB, where OECs lie adjacent to OSN axons. Accordingly, under the same experimental conditions that we observed weak inwardly rectifying currents in a sub-population of OECs from the rat OM, the currents in OECs derived from the rat OB did not show weak rectification. In common with previous observations from some OECs in the mouse OB (Rela et al., 2015), the Ba^2+^-sensitive currents found here in the majority (18/25) of OECs from the rat OB displayed strong inward rectification, suggesting a contribution by Kir channels with different properties to those of Kir4.1. The molecular identity of these channels remains undetermined. In this majority group of cells, there were also Ba^2+^-insensitive outwardly rectifying currents. We did not investigate the basis of any heterogeneity in the membrane properties of rat central OECs, which has been described extensively elsewhere (Rela et al., 2010; Rela et al., 2015). The biological importance of this heterogeneity remains to be determined.

Large Kir4.1+ cells were evident within the deeper layers where OSNs relay with projection neurons. Although we did not confirm their identity, these may be astrocytes known to extend their processes into the synaptic regions within the glomeruli (Raisman, 2001). The continuous Kir4.1 labelling within the OM, but not within the ONL, suggests that Kir4.1 subunits contribute only to OEC function in the periphery. Indeed, the distinct regional properties of the OECs recorded in this study, which reflect the differences in their expression of K^+^ channel subunits, point to fundamentally contrasting physiological roles for glia in the olfactory periphery and the brain.

The homeostasis and regeneration of neurons often relies on a bi-directional communication that informs the attendant glia of ongoing neural activity. Studies using OB slice preparations have revealed glutamatergic and purinergic receptor-mediated signaling between OSN axons and OECs (Rieger et al., 2007). This metabotropic mechanism involves a contribution from intracellular Ca^2+^ stores (Hayat et al., 2003), Ca^2+^ influx channels (Davies, Hayat, Wigley, & Robbins, 2004), and intracellular Ca^2+^ waves mediated by gap junctions (Stavermann et al., 2015). There is evidence of several connexins expressed by the non-neuronal cells in the OB (Rash et al., 2005), and genetic deletion of Cx43 leads to the loss of intercellular gap junctional coupling between OECs (Piantanida et al., 2019). There is less known about the physiology of peripheral OECs. Our whole-cell recordings have now revealed potential roles for weakly rectifying K^+^ channels in the normal function of the olfactory periphery. A consequence of this finding is that we might now consider Kir4.1 as a functional marker of OECs within this tissue, both in experimental animal studies and within tissues excised from humans. It remains to be determined though, whether Kir4.1 expression is detectable within the intact human OM, as our experiments here did not clearly demonstrate it. If Kir4.1 does contribute to normal homeostasis of OSNs, this may have consequences for our understanding of certain forms of anosmia, and of genetic conditions in humans such as EAST syndrome (Bockenhauer et al., 2009), in which sensory deficits result from mutations of the gene coding for Kir4.1 protein.

The continual turnover of OSNs during normal olfactory function demonstrates the potential for OECs to repair axonal damage in the human CNS. The answer to how OECs promote neuronal regeneration and regrowth of their axons may be multi-factorial. There is evidence for several mechanisms mediated by transplanted peripheral and central OECs. These include neuroprotection and increased axonal sprouting (Li, Field, & Raisman, 1997; Novikova, Lobov, Wiberg, & Novikov, 2011; Ramon-Cueto, Plant, Avila, & Bunge, 1998; Richter, Fletcher, Liu, Tetzlaff, & Roskams, 2005), recruitment of other beneficial cell types (Richter et al., 2005), remyelination (Franklin, Gilson, Franceschini, & Barnett, 1996; Imaizumi, Lankford, Waxman, Greer, & Kocsis, 1998; Radtke et al., 2004), and modification of the glial scar (Ramer et al., 2004). Due to the lower risk and invasiveness of the procedure, some researchers prefer biopsy of OECs from the OM rather than from the OB (Choi & Gladwin, 2015; Yao et al., 2018). These cells may have additional advantages brought about by their increased proliferative abilities (Jani & Raisman, 2004), and a greater capacity for migration and axonal growth (Richter et al., 2005). Further comparative studies of OECs from the OM and OB will be required to unravel the physiological differences between them, and to explain potential variability in their regenerative capacity.

Despite numerous reports of success following OEC transplantation into the CNS, there are also a number of studies with variable or negative outcomes. These may be a consequence of the discrepancies in the experiments and methods of assessment. The site of origin, the composition of cultures and their proliferative state may also be crucial factors that determine success following transplantation, and these have been reviewed recently elsewhere (Assinck et al., 2017; Gladwin & Choi, 2015; Gomez et al., 2018). The regenerative capacity of OECs may be determined by the length of time that they are cultured prior to transplantation. OECs cultured for short periods were more successful in terms of neuroprotection and growth sprouting than those maintained for longer *in vitro* before transplantation (Novikova et al., 2011). Our results suggest that OECs cultured from both rat and human OM express weak rectifier channels following their excision from the olfactory periphery. This observation may be relevant to the question of their proliferative state in culture. Pharmacological block of weak inwardly rectifying currents in quiescent astrocytes causes increased proliferation, and results in an increased proportion of cells in S-phase (Higashimori & Sontheimer, 2007). Ion channels expressed in cultured OECs could determine their proliferative state, and thus determine their likelihood of achieving both the growth in cell numbers and the de-differentiation required to maximize regeneration after transplantation.

Our work has demonstrated a functional distinction between the OECs of the olfactory periphery and those within the brain. It has indirectly identified the Kir4.1 channel subunit as a functional biomarker for peripheral OECs, potentially enabling their identification within OM endoscopic biopsies where numerous cell types are unavoidably present. This finding may help refine methods for their purification. Our study also verifies the adult rat OM as an experimental model for characterizing the physiology of human OECs. Further studies utilizing more intact preparations will be required to delineate the complex physiological roles of OECs within the normal olfactory system, and their interactions with neuronal axons in the periphery. Subsequently, we may gain a better understanding of how the membrane proteins underlying these normal processes change their expression following excision of OECs from the nasal mucosa. By targeting specific ion channel mechanisms and manipulating the membrane physiology of cultured OECs, we may be able to modulate the regenerative capacity of these cells for transplantation therapies.

## Acknowledgements

This study was supported by an Action on Hearing Loss/Dunhill Medical Trust Pauline Ashley Fellowship to KES (PA24), grants from the Biotechnology and Biological Sciences Research Council to DJJ (BB/M019322/1 and BB/R000549/1), and a grant from the Wellcome Trust to DC (200134/Z/15/Z). The work was carried out in part at the UCL/UCLH Biomedical Research Centre.

